# The Impact of Hand Movement Velocity on Cognitive Conflict Processing in a 3D Object Selection Task in Virtual Reality

**DOI:** 10.1101/2020.04.21.053512

**Authors:** Avinash K Singh, Klaus Gramann, Hsiang-Ting Chen, Chin-Teng Lin

**Author notes:** Corresponding author. Australian Artificial Intelligence Institute, School of Computer Science, Faculty of Engineering and Information Technology, University of Technology Sydney, Ultimo 2007, NSW, Australia. E-mail address (Avinash K Singh).

## Abstract

Detecting and correcting incorrect body movements is an essential part of everyday interaction with one’s environment. The human brain provides a monitoring system that constantly controls and adjusts our actions according to our surroundings. However, when our brain’s predictions about a planned action do not match the sensory inputs resulting from that action, cognitive conflict occurs. Much is known about cognitive conflict in 1D/2D environments; however, less is known about the role of movement characteristics associated with cognitive conflict in 3D environment. Hence, we devised an object selection task in a virtual reality (VR) environment to test how the velocity of hand movements impacts human brain responses. From a series of analyses of EEG recordings synchronized with motion capture, we found that the velocity of the participants’ hand movements modulated the brain’s response to proprioceptive feedback during the task and induced a prediction error negativity (PEN). Additionally, the PEN originates in the anterior cingulate cortex and is itself modulated by the ballistic phase of the hand’s movement. These findings suggest that velocity is an essential component of integrating hand movements with visual and proprioceptive information during interactions with real and virtual objects.

## Introduction

Several mechanisms are involved when humans interact with their environment, each making use of information from different sensing modalities, such as visual cues and proprioception(Scheidt, Conditt, Secco, & Mussa-Ivaldi, 2005). These sensory modalities serve the brain’s monitoring system, which instructs, plans, and executes interactions (Ozkan & Pezzetta, 2018). Importantly, this monitoring is constant to ensure one’s perceptions of their surroundings are continually updated (Singh et al., 2018) to match reality (Padrao, Gonzalez-Franco, Sanchez-Vives, Slater, & Rodriguez-Fornells, 2016; Padrao, Rodriguez-Herreros, Perez Zapata, & Rodriguez-Fornells, 2015). Should a change occur ‘mid-strategy’, i.e., during the process of planning and executing an interaction, the result is a mismatch response known as cognitive conflict (Fan, Flombaum, McCandliss, Thomas, & Posner, 2003). The human brain makes predictions about the outcome of an interaction, continuously comparing perceived information to that prediction, and when the prediction fails to hold, conflict occurs.

Cognitive conflict was first discussed in an article by Donchin et al. (1988) and republished in Donchin and Coles (2010). While there was no specific mention of event-related potential (ERP) related to an error, the work of Donchin and colleagues described P300 amplitude modulations due to changes in the environment. Later, Coull and Nobre (1998) showed that cognitive conflict causes one to redirect attention and reconfigure their initial plan, causing higher cognitive resources than non-conflict. In subsequent years, a first systematic experimental task was devised for cognitive conflict, known as the bimanual choice reaction task, which revealed that cognitive conflict causes a sequence of two types of ERP. First, the erroneous response causes an error-related negativity (ERN or Ne), which is a negative ERP typically peaking at around 50–150 ms (Falkenstein, Hohnsbein, Hoormann, & Blanke, 1991; Gehring, Goss, Coles, Meyer, & Donchin, 1993). This is followed by error-related positivity (Pe) after the erroneous response begins, which typically peaks at around 200–400 ms. Since this discovery, several experimental scenarios have been developed to test and demonstrate ERN and Pe. These scenarios include tasks like the Eriksen flanker task (Eriksen & Eriksen, 1974; Kopp, Rist, & Mattler, 1996), the oddball task (Halgren, Marinkovic, & Chauvel, 1998; Squires, Squires, & Hillyard, 1975), and the Stroop task (Stroop, 1935; West & Alain, 1999). Some other variants of ERN include feedback-related negativity (FRN) (Holroyd & Coles, 2002), and observational error, due to a person observing another person making an error (van Schie, Mars, Coles, & Bekkering, 2004).

However, most of the experiments, protocols, and findings described above only pertain to passive one-dimensional (1D) or two-dimensional (2D) stimuli and cannot necessarily be generalized to a three-dimensional (3D) world. A realistic 3D input, for example, grasping a bottle on a table in front of you, adds significant complexity to an interaction task given the computations need to move one’s body parts through space.

At the same time, realistic 3D interactions provide a window of opportunity to understand better the brain’s monitoring function and how it conducts complex monitoring of the real world (Jungnickel & Gramann, 2016). One of the most basic 3D interactions is an object selection task (Ferran Argelaguet & Andujar, 2013). To grasp an object in the 3D world using the hand, the user is required to perform a set of complex movements that involve positioning their palm and fingers over the object. Our previous work with a 3D object selection task demonstrated that changing the selection radius of a virtual cube (leading to premature feedback of touching the object) could lead to a mismatch between the visual and the proprioceptive feedback, invoking cognitive conflict (Gehrke et al., 2019; A. K. Singh, H.-T. Chen, K. Gramann, & C.-T. Lin, 2020; A. K. Singh, H. Chen, K. Gramann, & C. Lin, 2020; Singh et al., 2018). We found the conflict reflected a form of prediction error negativity (PEN) that seems to belong to the same class of ERP as the ERN and FRN found in the studies mentioned above. Additionally, the results indicated that sensory integration plays an essential role in monitoring and producing ongoing actions, particularly for 3D object selection. Our findings also suggest that visual feedback dominated proprioceptive feedback in participants who completed the task in a short amount of time. We thus concluded that proprioceptive feedback is more important for slower movements. However, limitations in the experimental protocol did not allow us to analyze hand movement velocity and its role in integrating visual and proprioceptive information.

To overcome the limitations of our previous studies and to further investigate the role of movement velocity on the electrocortical responses to cognitive conflict, we designed an experimental protocol that manipulated hand movement velocity. Our intuition was that the velocity at which a hand moves does impact cognitive conflict processing in 3D object selection tasks. To test this hypothesis, we devised a task to measure hand movement velocity in tandem with brain electrical responses to cognitive conflict using two different sizes of the cube to invoke two kinds of hand movement velocity profiles. By manipulating the hand movement velocity, we manipulated the hand and arm movement of participants in different experimental conditions leading to differences in proprioceptive feedback. The concurrent processing of proprioceptive feedback and its comparison with the predicted feedback when an action is executed, is one major sensory source for controlling the accuracy of the movement itself (Desmurget & Grafton, 2000). Deviations of the proprioceptive feedback from the predicted movement feedback should thus contribute to error detection and the associated brain dynamic markers like the PEN.

## Materials and methods

### Participants

To determine the effect of movement velocity on the amplitude of PEN, we recorded electroencephalogram (EEG) data from 20 participants (2 females and 18 males) with all right-hand dominated except one participant. The mean age of the participants was 23.3 years, with a range of 18-30 years. Before participating in the study, each participant was given a full explanation of the experimental procedure, and each provided informed consent. Ethics approval was issued by the Human Research Ethics Committee of the University of Technology Sydney, Australia. The experiment was conducted in a temperature-controlled room by a male experimenter. None of the participants had a history of neurological or psychological disorders, which could have affected the experiment results; participants were allowed to wear glasses for corrected vision.

The number of participants took part in this study have a large effect from the velocity of the hand’s movement on the amplitude of PEN (F (1, 17) = 89.454, p < .001, η2 = 0.99) based on the calculation of Green, Salkind, and Akey (1997).

### VR setup

The virtual reality (VR) environment was provided through an HTC Vive head-mounted OLED display with a resolution of 2160 × 1200 and a refresh rate of 90 Hz (HTC Corp., Taiwan). The participants’ head positions were tracked with the embedded inertial measurement units (IMUs), while an external Lighthouse tracking system cleared the common tracking drift with a 60 Hz update rate.

Hand motions were recorded with a Leap Motion controller (Leap Motion Inc., USA) attached to the front of the HTC Vive that tracked the fingers, palms, and arms of both hands up to approximately 60 cm above the device with 120 frames/second. The tracking accuracy has been reported to be 0.2 mm (Weichert, Bachmann, Rudak, & Fisseler, 2013) and the latency has been reported to be approximately 30 milliseconds (Bedikian, 2013).

### EEG setup

The EEG data were recorded from passive 64 Ag/AgCl electrodes, which were referenced to an electrode placed between locations Cz and CPz. The placement of the EEG electrodes was consistent with the extended 10% system (Chatrian, Lettich, & Nelson, 1985). Contact impedance was maintained below 5kΩ. The EEG recordings were collected using a Curry 8 SynAmps2 Express system (Compumedics Ltd., VIC, Australia) with a digital sample rate of 1 kHz in 16-bit resolution.

First, the participants were equipped with an EEG cap and an additional separator cap (a plastic shower cap) to reduce electrolytes polluting the VR equipment. The head-mounted display was placed on top of the separator cap (see Figure 1). To ensure the participants had a better VR experience, we installed a table similar to the one used in the VR environment. The height of the table in both worlds was similar in color, and height, so participants were not able to distinguish between the real and the virtual environment.

**Figure 1.**
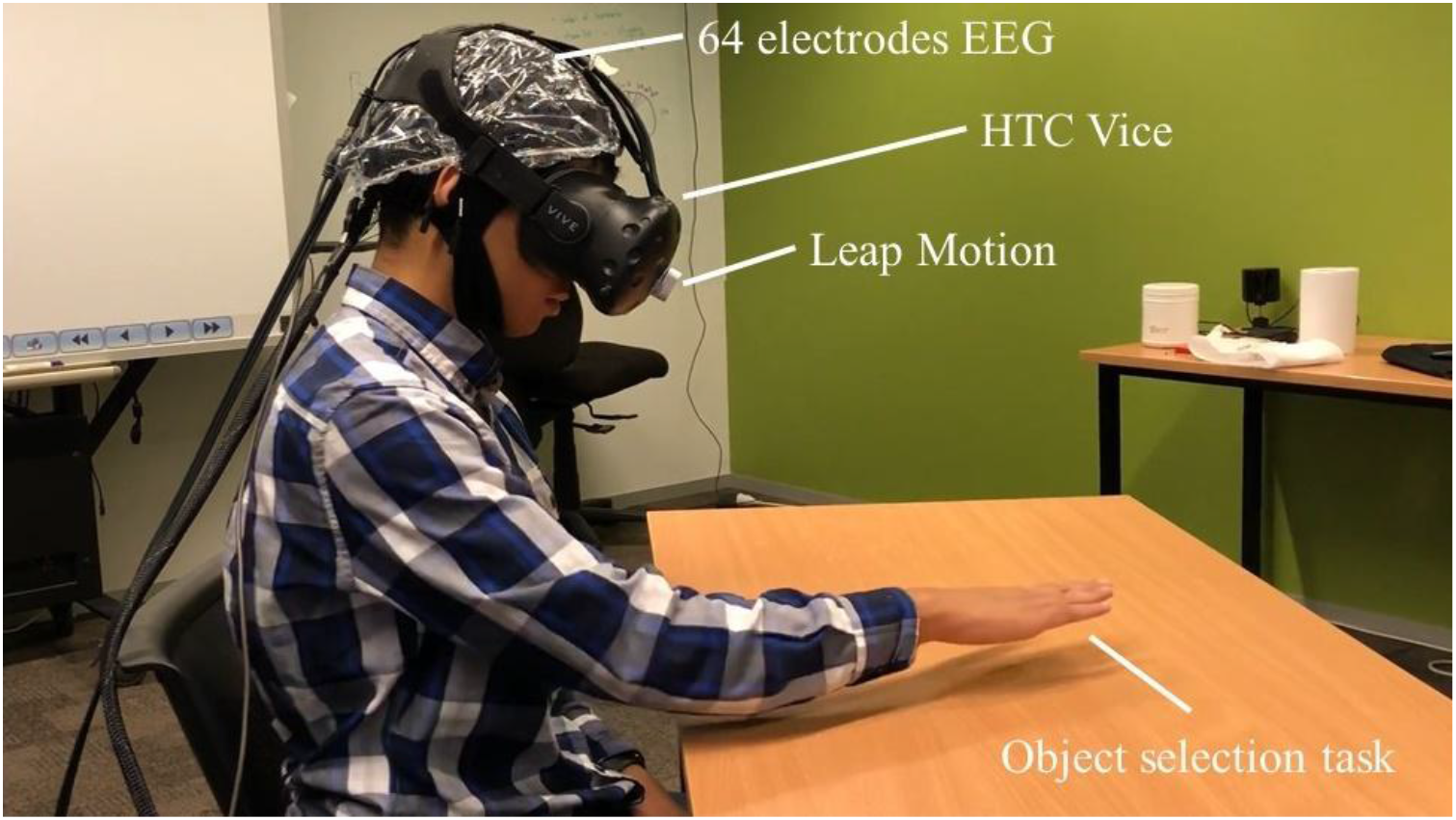
A participant performs the 3D object selection task with an HTC Vive head-mounted display and a Leap Motion controller while wearing the 64-electrode EEG cap

### The experiment scenario

Each participant performed the 3D object selection task with their dominant hand tracked. Figure 2 displays a scenario for a single trial. Each trial was four seconds long. The scenario started with instructions about the task, followed by the experimental trials. In each trial, a cube appeared on the table, which the participants were instructed to reach out to and touch. The cube turned red when touched as a feedback signal. Participants were expected to finish the task within 4 seconds; otherwise, the trial was marked as incomplete.

**Figure 2.**
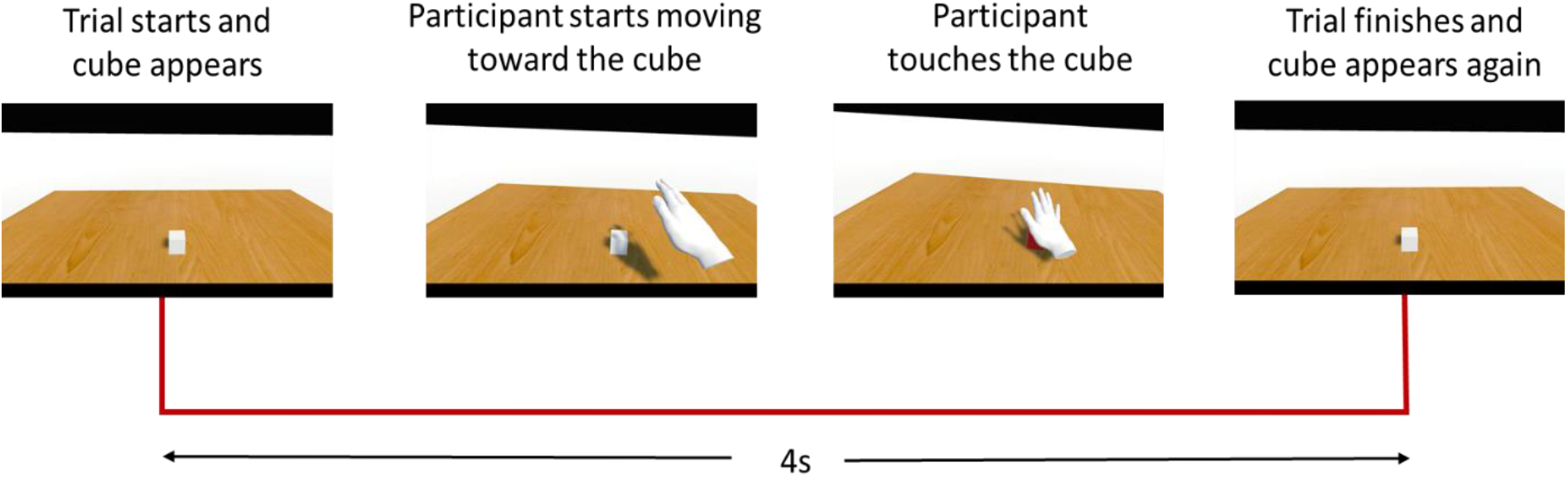
Experiment scenario for a single trial

The experiment was designed with two degrees of difficulty – a small cube (*d*) and a big cube (*D*) – which each produce a distinct velocity profile based on Fitts’s law (Soukoreff & MacKenzie, 2004). Selecting a small cube was more difficult than selecting a big cube as the endpoint of the movement required more finely-grained motor adjustments resulting in a lower velocity.

The cognitive conflict condition was invoked by manipulating the selection radius on the cube. The selection radius is the radius of an invisible sphere surrounded the cube that is used in the virtual reality environment to measure the interaction of other objects or agents in the VR with this object. Once the system detects the virtual hand is in contact with the invisible sphere (to select the object), the cube changed its color from white to red. There were two kinds of selection radii for the non-conflict and the conflict conditions. The selection radius of the invisible sphere equaled the size of the cube, i.e., ‘d’ for the non-conflict condition. In this case, the color change indicating that the objects were touched appeared at the moment when the hand of the participants touched the virtual object. In the conflict condition, in contrast, the selection radius was 1.5x larger than the radius ‘d’ leading to a color change of the object before the hand reached the actual object. The participant naturally expected the cube to change its color when the virtual hand reached it, i.e., at a distance ‘d. In the conflict condition, the cube changes its color prematurely when the hand reached the cube, i.e., at distance D = 1.5d. See Figure 3 (A).

**Figure 3.**
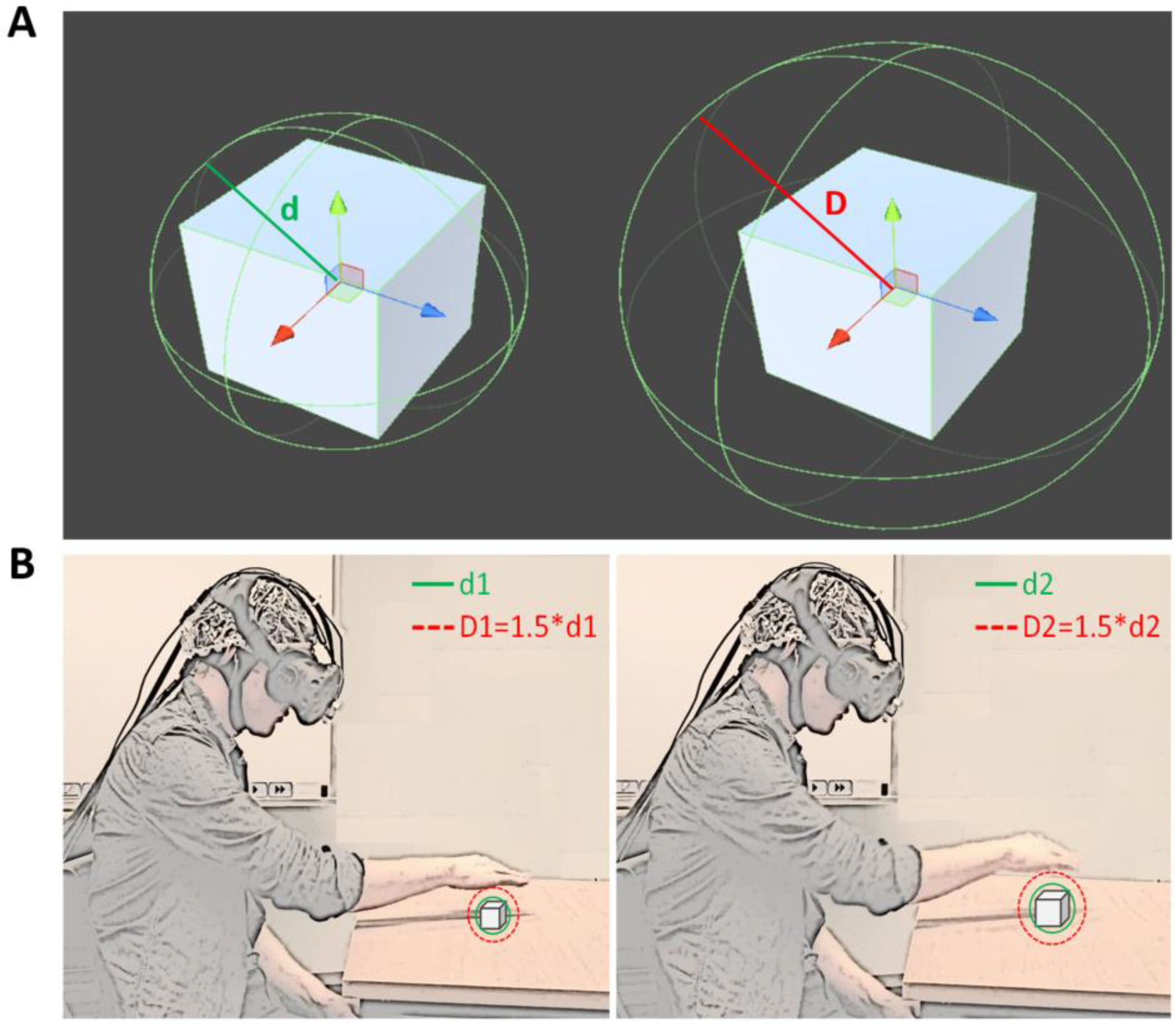
A. a cube representation from Unity^2^ software surrounded by a sphere (invisible) of green (d) or red radius (D) at a time based on condition, B. a schematic representation of the 3D object selection task performed by a participant with small (left) and big cubes (right) with non-conflict (d1 and d2) and conflict (D1 and D2) selection radius. The selection radius is not visible to participant.

The selection radius of the cubes, defining the collision between the endpoint of the moving hand to the cube, changed in 25% of the trials, such that 75% of the trials used distance D1/d1 and the remaining trials used distance D2/d2 for big/small cubes (D1=1.5*d1 and D2=1.5*d2; See Figure 3(B)). The cube was also placed in three positions: in the center (0°), at 30° to the left, or at the same radius at 30° to the right. This created variety in the velocity profiles and kept the participants engaged.

The experiment used a 2 by 2 design with two independent variables: a) the conditions – no-conflict and conflict; and b) the size of cubes – small and big. The experiment was conducted over two blocks with each block comprising 250 trials with an overall duration of about ~16 minutes. The full experiment for each participant took about 1.5 hours, including the initial setup of the EEG cap and head-mounted display, the trials, and completing the questionnaire.

### Questionnaire

While there is no standard method for measuring the presence in immersive virtual environments, most researchers use questionnaires to assess self-reports from users. In the current experiment, a modified version of Igroup presence questionnaire, i.e., IPQ (Schubert, 2003), together with the participant’s experience of game playing, was used to measure the presence of participants. The questionnaire comprised a total of 24 questions to be answered on a seven-point Likert scale. At the end of the experiment, each participant completed a 24-item IPQ asking them to rate different parameters of the experiment on a 7-point Likert scale, such as realism, experience, and controlling events, to yield a possible result of between 7 and 98. Additionally, the IPQ included a space asking them to state their previous experiences with game playing to help assess their overall proficiency with VR scenarios.

## Data Analysis

### EEG data analysis

We used the EEGLAB toolbox (Delorme & Makeig, 2004) in MATLAB 2016 (MathWorks Inc, USA) to process the EEG data. The raw EEG signals were filtered using a 0.1-Hz high-pass and 40-Hz low-pass FIR filter with a filter order of 15 with zero-phase and subsequently downsampled to 250 Hz. The resulting data were inspected to identify and remove noisy electrodes using the Kurtosis method, followed by an ICA (Makeig, Bell, Jung, & Sejnowski, 1996) and equivalent dipole model fitting using DIPFIT with a spherical four-shell (BESA) head model (Scherg, 1990). The resultant ICs were further processed to detect artifact-related ICs using the SASICA plugin (Chaumon, Bishop, & Busch, 2015), which uses autocorrelation, focal ICs, eye blinks, and information from the ADJUST plugin (Mognon, Jovicich, Bruzzone, & Buiatti, 2011) to identify ICs representing artifacts. These ICs were marked and excluded from the final data. On average, 19.80 ± 8.82 ICs were removed, and the data was back-projected to the sensor level. The back-projected data were epoched from 500 ms prior to touching the cube to 1000 ms after the touch event for all conditions, as well as being inspected again for artifacts using the Kurtosis method. On average, 19.84 ± 10.04% epochs were removed. Please see Supplementary Figure 1 for example of exemplary detected ICs.

The IC components of all the participants were clustered using a neural network-based clustering approach, implemented in EEGLab, based on their similarity with respect to the ERP, power spectrum, and event-related spectral perturbations (ERSP), plus the component scalp maps, their equivalent dipole, and corresponding dipole locations for each participant. The approach was able to cluster the IC components shared by approximately 70% of the participants. Our more in-depth analysis focuses on clusters with IC components located in or near the cingulate cortex (Montgomery, Huang, & Assadi, 2005) to find cognitive conflict (Schlüter et al., 2018) related IC. The clustered components were further used to compute ERSP and to extract the PEN and Pe from the back-projected and channel-based ERPs. PEN was calculated as the minimum amplitude in a search window of 50-150ms after the touch event calculated as the mean of the minimum ±2 adjacent sample points. Similarly, Pe was calculated as the mean of the maximum in a 250-350ms search window after the touch event, including ±2 adjacent sample points.

### Behavioural data analysis

#### Task completion time

Task completion time was calculated as the difference between the time the cube appeared until the cube changed color.

#### Hand motion trajectory data

Velocity, hand position (palm, finger, etc.), the number of frames in each VR scene, and the time required for each frame were recorded for each trial for each participant. The primary focus of this experiment was to understand how the velocity profile of the reaching movement affected the PEN. Therefore, the velocity and acceleration were calculated for each trial at each sample point during the hand movement.

#### Extraction of a ballistic and corrective phase of velocity

The hand movement velocity was divided into two parts based on the maximum peak of the velocity of each trial. The velocity before the peak is known as the ballistic phase of the hand-movement velocity. The velocity after the peak is known as the corrective phase of the hand-movement (Meyer, Abrams, Kornblum, Wright, & Smith, 1988). Therefore, designated the initial movement phase before the peak velocity as the ballistic phase and the remaining movement as the corrective phase. To further concentrate on the most representative ballistic and corrective phase, we further selected the first 20% of the segmented ballistic data for further analysis, and similarly, the last 20% of the corrective phase of velocity data as the absolute corrective phase of hand movement for further analysis.

#### Statistical analysis

All the statistical analyses of behavioral and EEG data were conducted using SPSS (IBM SPSS Inc Version 24) and Matlab 2016 (Mathworks Inc, USA).

##### ANOVA

A 2 × 2 repeated analysis of variance (ANOVA) was conducted using SPSS for task completion time with the factors a cube sizes (small vs. big cube) and conflict conditions (non-conflict vs. conflict) with a significance level of alpha = .05 followed by post-hoc analysis using the Tukey method to determine the source of significant difference with .05 significance level if any.

T-test. One-sample t-tests were conducted using SPSS on difference PEN and Pe amplitudes between small and big cubes with a test-value of 0 for all 64 electrodes. To minimize the Type-1 error, 1000-fold permutation testing was applied. If the returned p-value was below .05 for a channel, it was marked as statistically significant.

Statistics for ERSP. Similar to ERP, difference ERSP between condition and cube size were investigated to find the time windows and frequency, revealing significant differences between small and big cubes for a trial. To define statistical significance between the unpaired values, the EEGLab function statcond() was used. Due to this, a 1000-fold permutation testing was applied. If the returned p-value was below .05, the sample was marked as statistically significant and plotted on a 2-d grid of frequency and time for each point.

##### ANCOVA

We have also conducted a 2 × 2 repeated measure analysis of covariance (ANCOVA) using SPSS with the factors a cube sizes (small vs. big cube) and conflict conditions (non-conflict vs. conflict) on PEN and Pe amplitude with task completion time as co-variate.

#### Regression analysis

The multiple linear regression was also conducted using SPSS (IBM SPSS Inc Version 24) for PEN and Pe w.r.t. the conflict/no-conflict condition. The entered variables were in the following order: the velocity of the hand’s movement during the ballistic phase, the VR quality score (QVR), the VR quality self-evaluation score (SVR), realism, the possibility of action in VR (PAVR) score, and the task completion time. We also performed another regression analysis swapping the order of the ballistics velocity with the task completion time. The results of the second regression analysis have been reported in the supplementary.

#### Correlation analysis

We tested the relationship between the ERSP for the delta, theta, alpha, and beta power bands and the ballistic phase of the hand movement velocity with a Pearson’s correlation analysis using Matlab 2016 (Mathwork Inc, USA).

## Results

### Behavioral results

Figure 4 shows the box plot for the average task completion time for all trials for all participants. A repeated-measures ANOVA was conducted to compare the task completion times for 2 (cube sizes) × 2 (conditions). There was a significant difference for both the conditions (F (1, 18) = 191.074, p < .001) and the cube sizes (F (1, 18) = 302.815, p < .001). The two main effects were qualified by their interaction cube sizes * conditions (F (1, 18) = 5.358, p = .032). Tukey post-hoc tests revealed no significant difference in task completion time (p = .178) in the non-conflict trials with the small and big cubes. There was also no significant difference (p = .085) in task completion time for the conflict and non-conflict trials. However, there was a significant difference (p < .001) between the task completion time for the small and big cubes.

**Figure 4.**
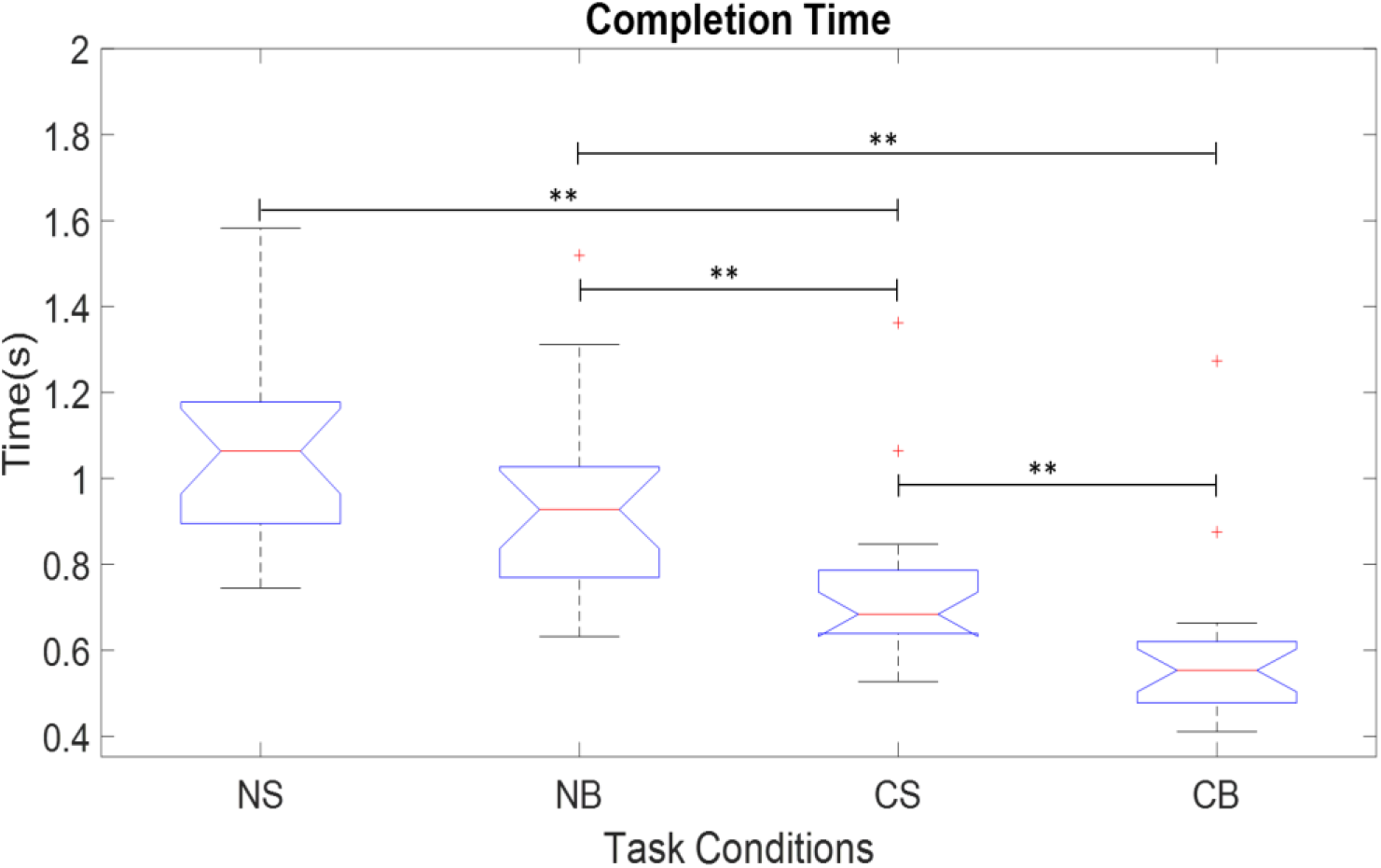
The task completion time for all conditions based on different selection radius for small and big cubes. NS) non-conflict trials with small cubes; NB) non-conflict trials with big cubes; CS) conflict trials with small cubes; CB) conflict trials with big cubes.

We also plotted the overall hand movement velocity pattern for all trials to see how it changed over time. Figure 5 (NS & NB) shows the hand’s trajectory initially increased sharply for about 20-25 frames, which is known as the ballistic phase, then slowly decreased until a cube was selected. According to the optimized initial pulse (OIP) model, this decrease is known as the corrective phase (Meyer et al., 1988). The trajectories for the small and big cubes in the non-conflict trials were quite similar. However, as can be seen from Figure 5 (CS & CB), with the small cubes, the participant steadily increased velocity until the ballistic phase then decreased pace during the corrective phase until the cube was selected. By contrast, with the big cubes, there was still a sharp increase in the ballistic phase, but the decrease in the corrective phase was very steady compared to the small cubes. This is likely because, with a big target, the participants were less precise with their hand movements in the ballistic phase, resulting in more corrective movements needed in the corrective phase, as opposed to the more careful approach from the outset with the smaller cubes.

**Figure 5.**
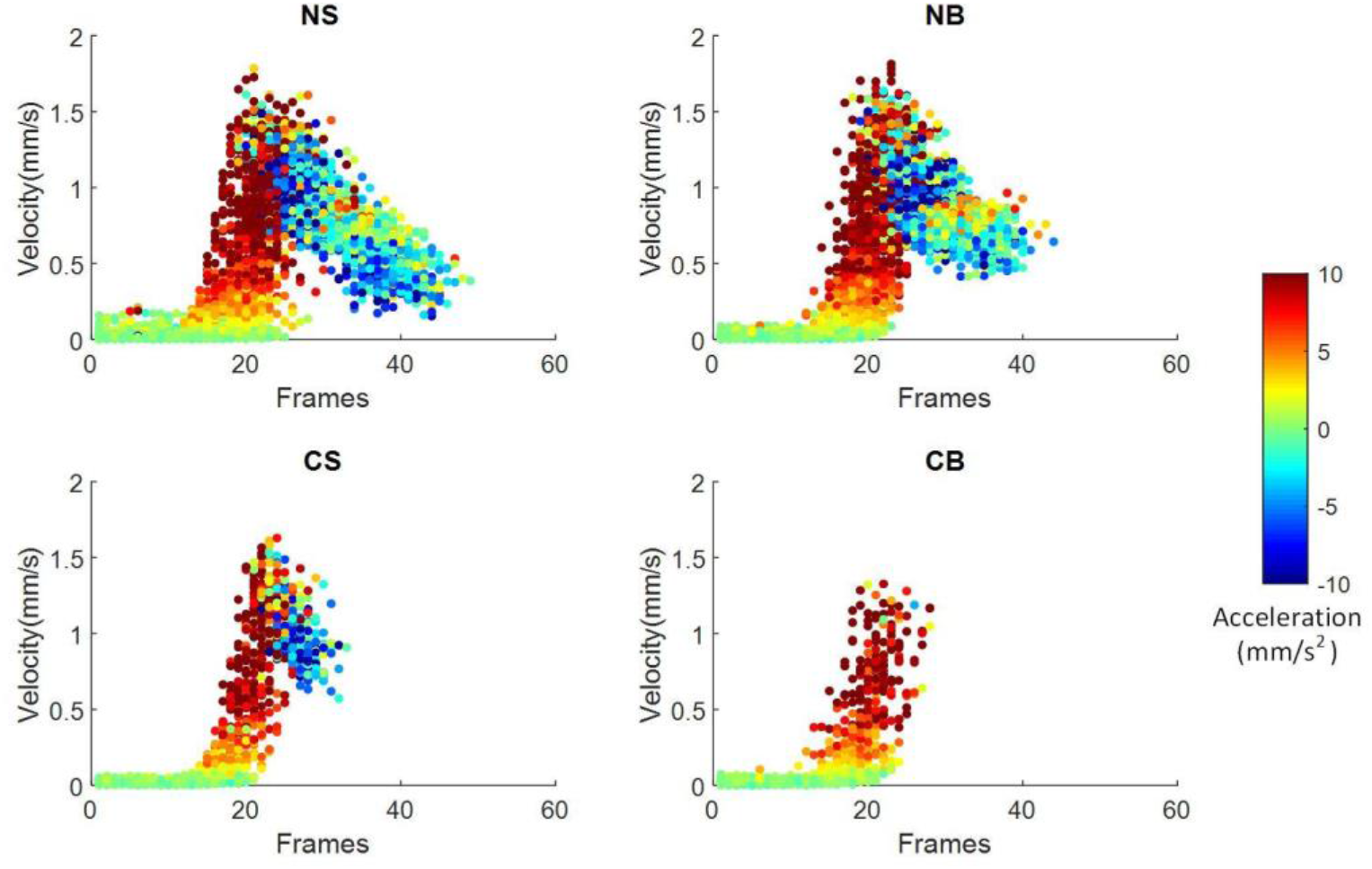
Hand movement velocities for all conditions for one exemplary participant. NS) non-conflict trials with small cubes; NB) non-conflict trials with big cubes; CS) conflict trials with small cubes; CB) conflict trials with big cubes

### EEG Results

Figures 6 and 7 show the topographical plots of the PEN and Pe, respectively. An independent samples t-test of the PEN component for both conditions indicate a significant difference in PEN at FCz in both the non-conflict and conflict trials for the small cubes (t (17) = −3.612, p = .002, p = .009) and FC2 in the big cubes (t (17) = −2.575, p = .020, p = .022). Similarly, there was a significant difference in Pe at channel FC6 in both the non-conflict and conflict trials with both the small cubes (t (17) = 2.178, p = .044, p = .077) and FCz for the big cubes (t (17) = −3.402, p = .003, p= .009).

**Figure 6.**
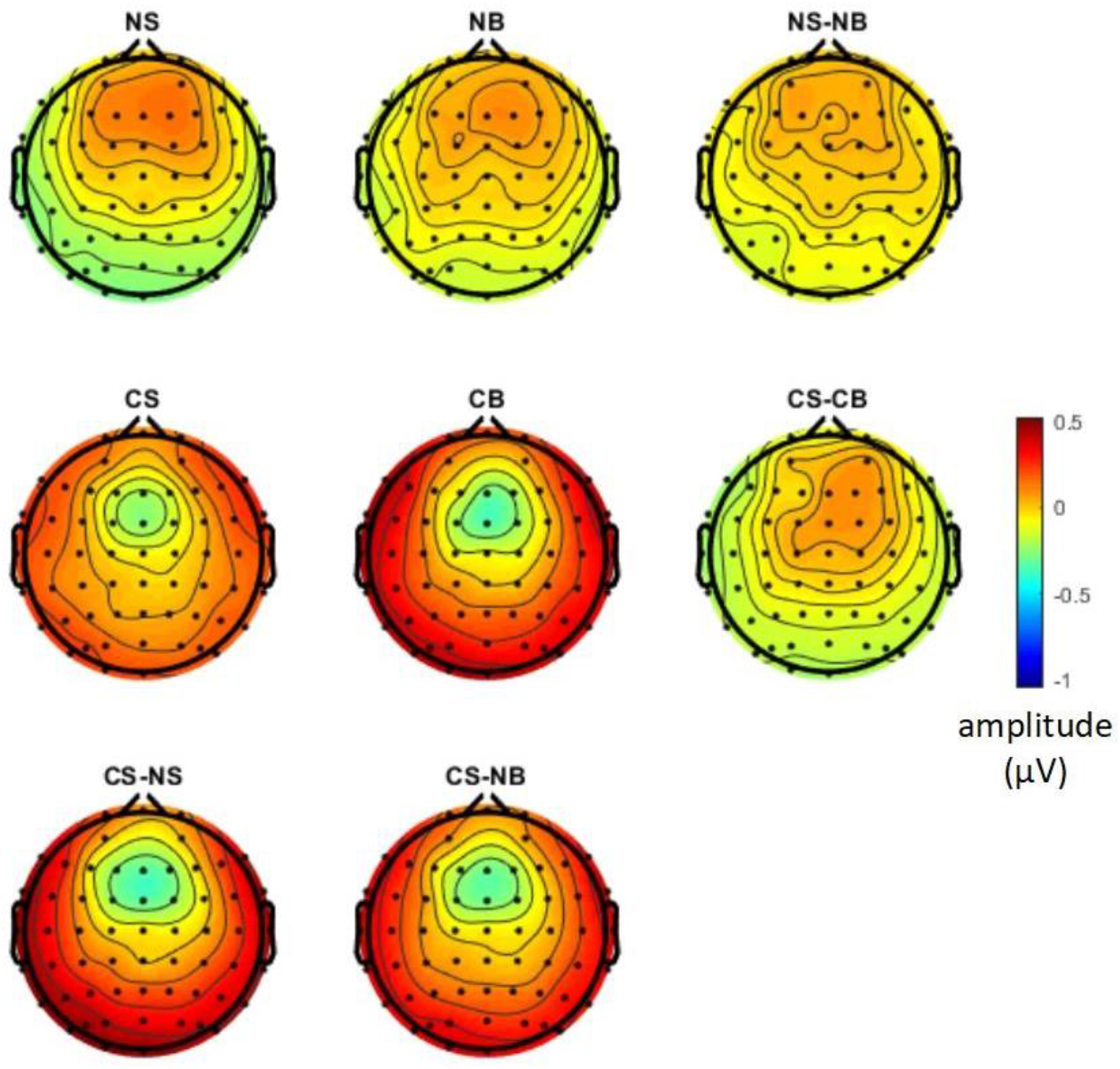
Topographical plots of PEN. NS) non-conflict trials with small cubes; NB) non-conflict trials with big cubes; CS) conflict trials with small cubes; CB) conflict trials with big cubes

**Figure 7.**
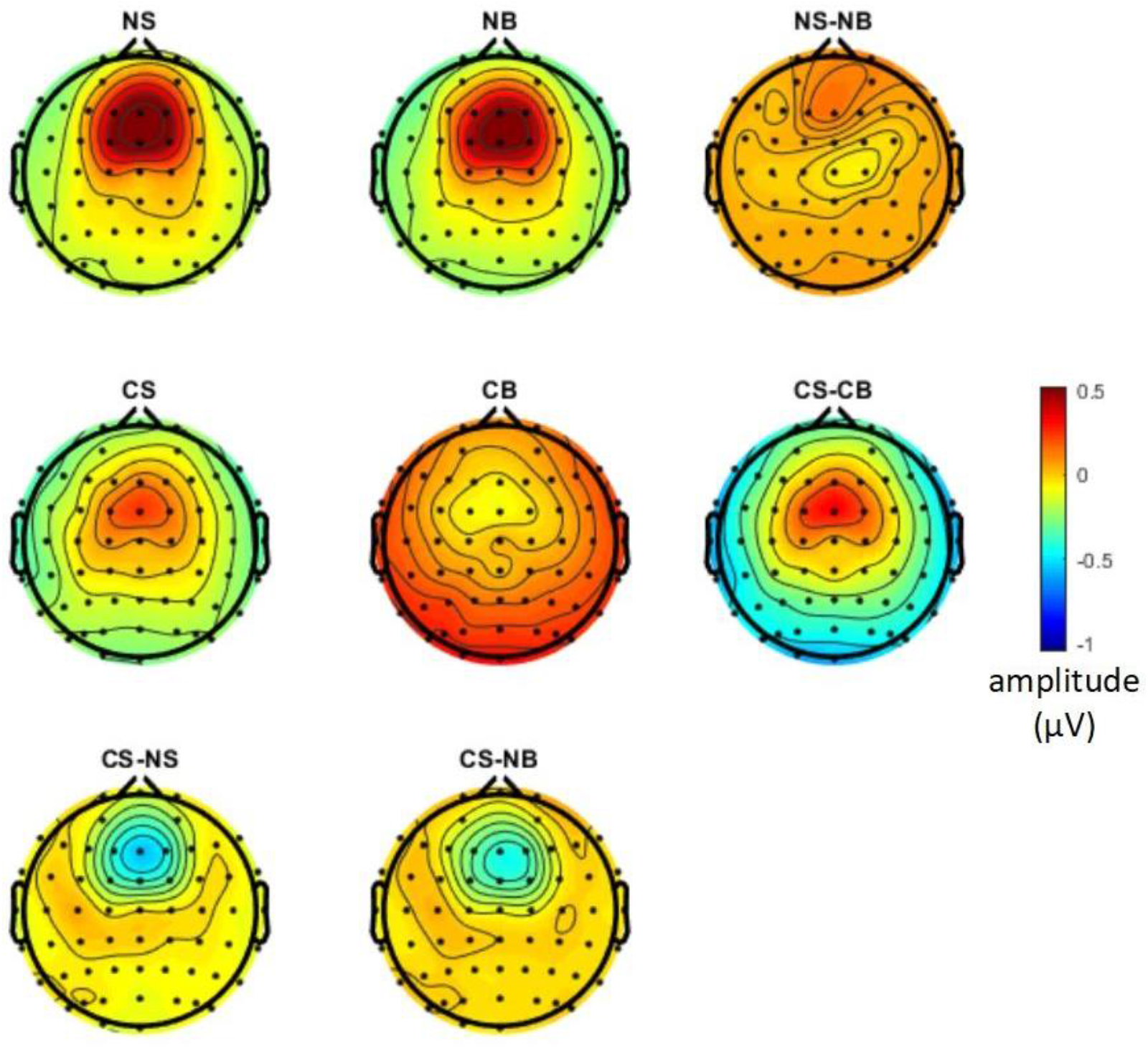
Topographical plots of Pe. NS) non-conflict trials with small cubes; NB) non-conflict trials with big cubes; CS) conflict trials with small cubes; CB) conflict trials with big cubes

As indicated from the topographical plots, the average ERP for all participants was evaluated for the fronto-central region at FCz. Figure 8 shows the difference ERP showing larger PEN for big cubes and larger Pe for small cubes.

**Figure 8.**
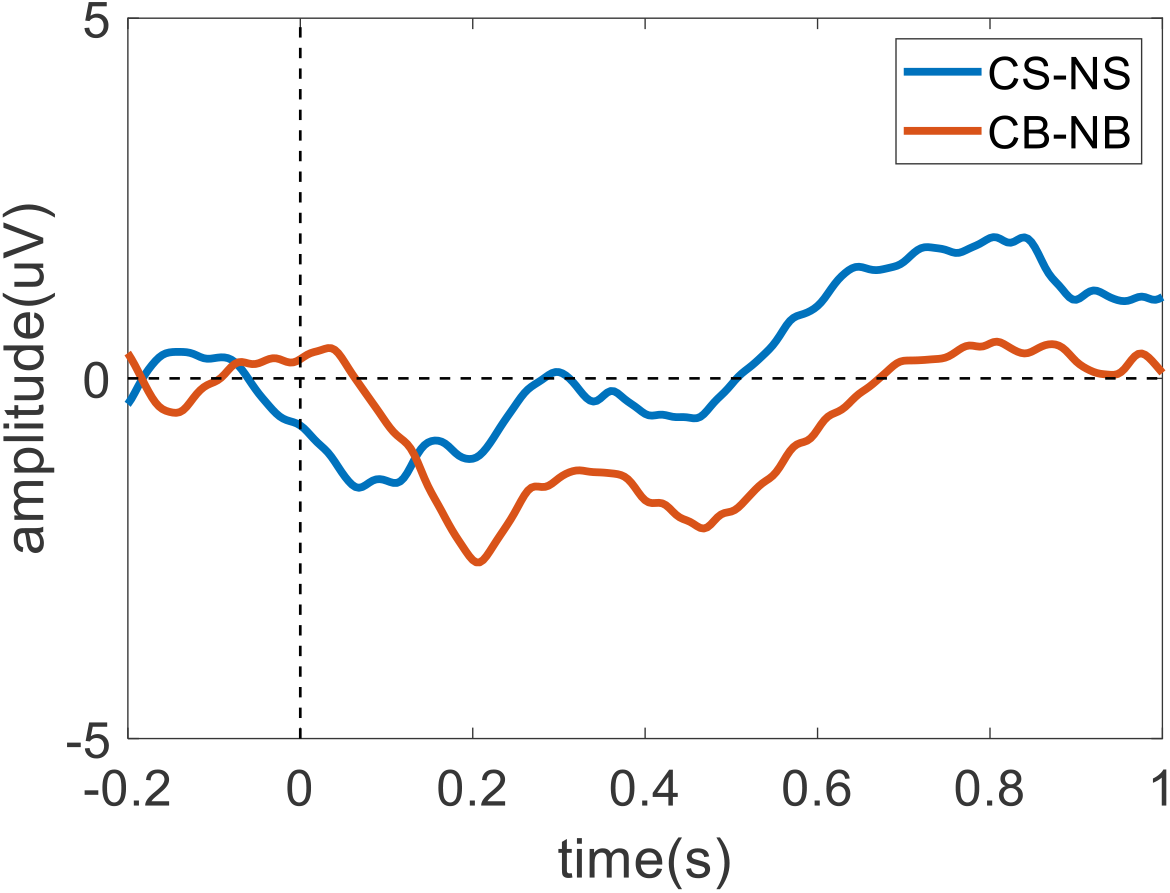
The difference in ERP between the no-conflict and conflict trials with the small and big cubes. NS) non-conflict trials with small cubes; NB) non-conflict trials with big cubes; CS) conflict trials with small cubes; CB) conflict trials with big cubes

By inspecting the properties of the different clustered ICs in the fronto-central region, we were able to extract the IC cluster with MNI coordinates of x=-6, y=10, and z= 24. According to the Talairach Client application^3^, MNI coordinates are identified as the anterior cingulate cortex (ACC) as shown in Figure 9. This component is shared by more than 70% of the participants.

**Figure 9.**
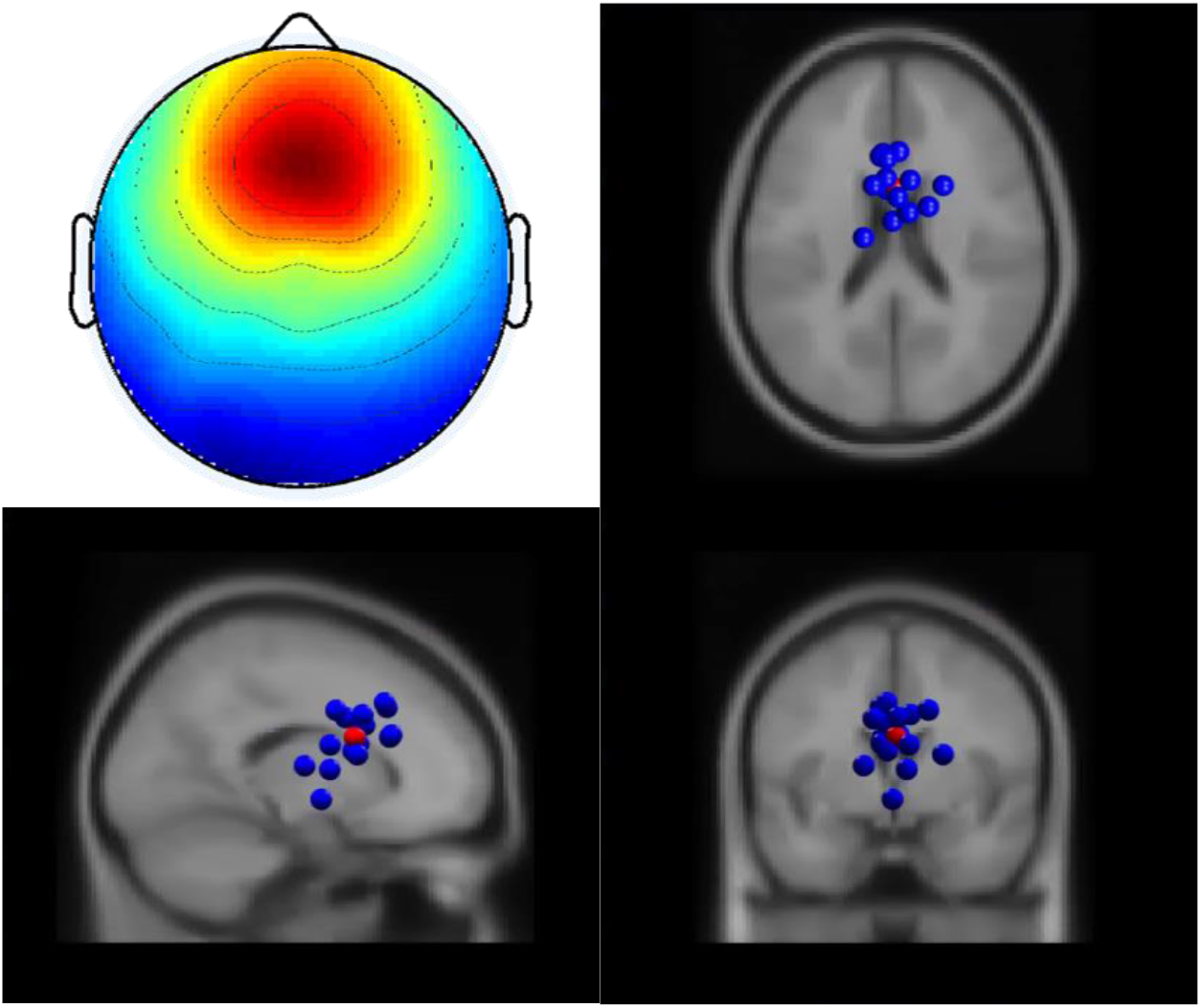
The identified anterior cingulate cortex and its dipole clusters from all participants (MNI coordinates x=-6, y=10, and z= 24)

We also found other clusters related to PEN. However, the results showed no difference between small and big cubes for PEN and Pe. See left and right-sensorimotor cortex based ICs and their ERP in Supplementary Figure 2.

To understand the effect of 3D object selection over time, we calculated the ERSP for the selected ACC component. As shown in Figure 10, in the conflict trials with the small cubes, there was substantial suppression in alpha band power between 50-150ms and in both the theta and alpha band at 400-700ms, which did not occur in the non-conflict trials. By contrast, in the conflict trials with the big cubes, there was suppression in beta band power at around 0-100ms and 300-600ms.

**Figure 10.**
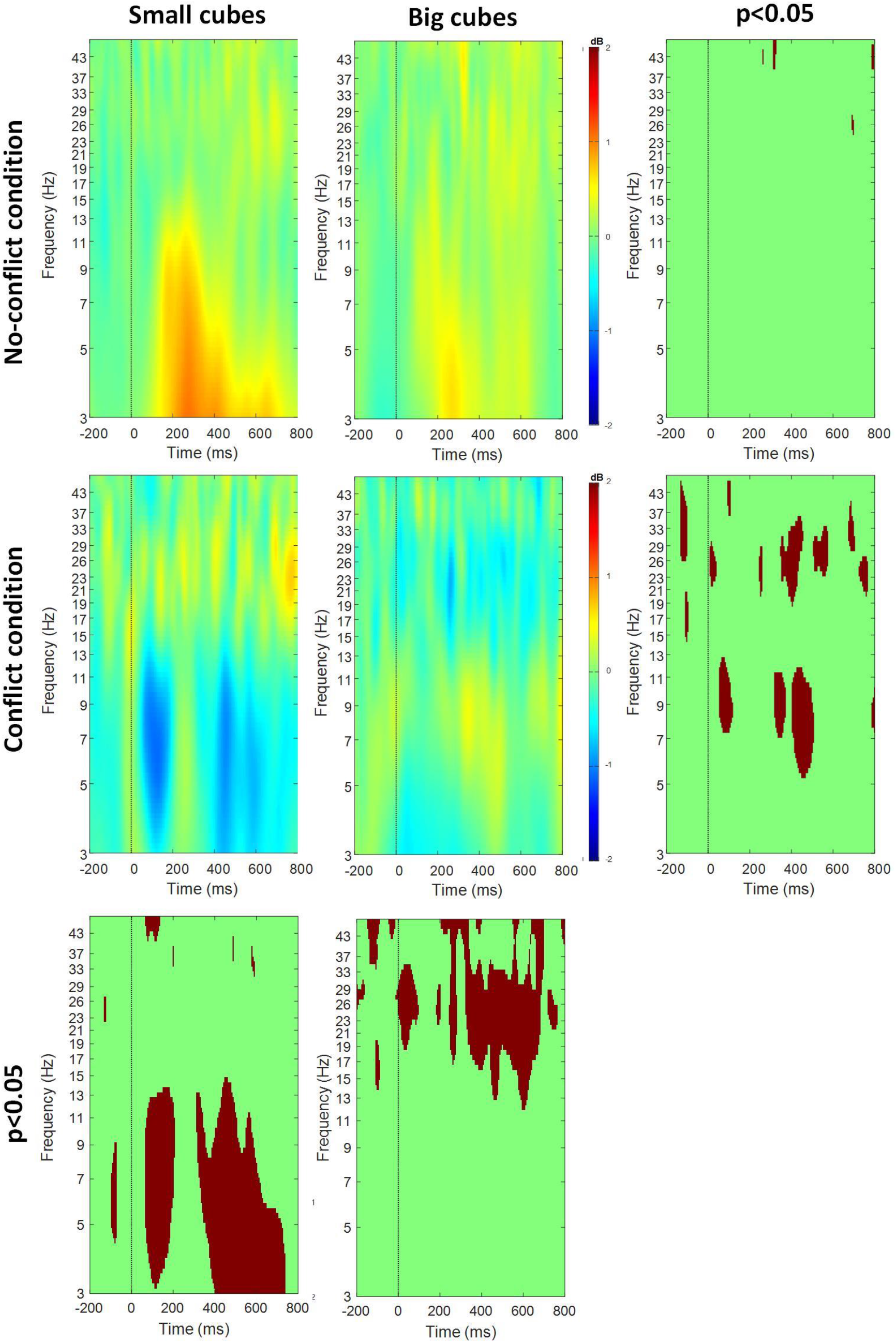
Event-related spectral perturbations and statistical results (brown color to represent significance at .05, otherwise green) for the big/small cube and conflict/no-conflict conditions

### Analysis of covariates

This analysis was to determine whether task completion time affected the PEN and Pe amplitudes for sizes × conditions design. Based on an ANCOVA with task completion time as the covariate, we found that task completion times do not affect the PENs. Therefore, there was a still a significant difference for both the conditions (F (1, 67) = 15.763, p < .001) and the cube sizes (F (1, 67) = 5.371, p < .001). Interestingly, there were no effect found by their interaction cube sizes * conditions (F (1, 67) = .472, p = .494).

However, task completion seems to have effect on Pe. There was no significant difference for both the conditions (F (1, 67) = .280, p < .599) and the cube sizes (F (1, 67) = .208, p < .650). Also, there were no effect found by their interaction cube sizes * conditions (F (1, 67) = 1.415, p = .238) either.

### Regression analysis

To understand the relationships between the subjective and behavioral data of the PEN and Pe amplitudes, we performed a regression analysis with the following results.

A linear regression established that the ballistic of hand movement velocity, together with the behavioral measures quality of virtual reality (QVR) and self-evaluation of virtual reality (SEVR), can predict PEN in conflict trials with small cubes (F (3, 17) = 4.455, p = .021). If added realism; the possibility to act in virtual reality (PAVR), task completion time, the ballistic of hand movement velocity, QVR and SEVR can also predict PEN in conflict trials with small cubes (F (6, 17) = 3.248, p = .043). However, we observed no such covariation for Pe in conflict trials with small cubes (F (6, 17) = .937, p = .507). The ballistic of hand movement velocity, together with QVR and SEVR, accounted for 48.8% of the variance in PEN amplitude. Including realism, PAVR, and task completion time raised that level to 63.9% of the variance. The regression equation to predict the PEN with small cubes was as follows:

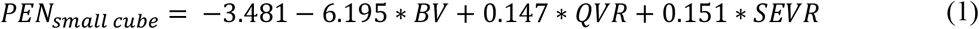

where BV = ballistic of hand movement velocity, QVR = quality of virtual reality scene, SEVR = self-evaluation of virtual reality scene.

Similarly, a linear regression established that the ballistic of hand movement velocity, together with the behavioral metrics taken from the IPQ, could predict the PEN amplitudes in the conflict trials with the big cubes at a statistically significant level (F (3, 17) = 3.880, p = .033). Again, when including realism, PAVR, task completion time, and the ballistic phase, QVR and SEVR could also generate an accurate prediction (F (6, 17) = 1.903, p = .043).

Notably, the ballistic of hand movement velocity alone was able to predict Pe amplitudes in conflict trials with big cubes at significant levels (F (1, 17) = 6.318, p = .023). The ballistic phase, together with the IPQ metrics, accounted for 45.4% variability in the PEN amplitude but only 28.3% for the Pe amplitude. The regression equation to predict the PEN with big cubes was as follows:

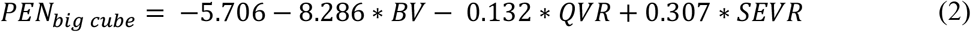

where BV = ballistic of hand movement velocity, QVR = quality of virtual reality scene, SEVR = virtual reality scene self-evaluation

The regression equation to predict Pe with big cubes was as follows:

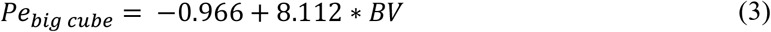

where BV = ballistic of hand movement velocity

### Correlation between velocity and spectral power

The Pearson’s correlation between the spectral powers (delta, theta, alpha, and beta) and the ballistic phase in the no-conflict trials with small cubes suggests that the ballistic phase was significantly correlated with the alpha band (r = − .200, n = 185, p = .006) and the beta band (r = −.181, n = 185, p = .014). However, we found no significant correlation for the conflict trials with small cubes. Yet with big cubes and the conflict trials, we found a statistically significantly correlation between the ballistic phase and the delta band (r = − .254, n = 61, p = .048) plus the theta band (r = − .323, n = 61, p = .011). No correlation was found in the non-conflict trials with big cubes.

## Discussion

We hypothesized that the velocity of the hand’s movement impacts cognitive conflict processing in 3D object selection and, more specifically, the amplitudes of the PEN and Pe components. To test this hypothesis, we designed a modified version of the 3D object selection task outlined in Singh et al. (2018) to produce two distinct hand movement velocity profiles based on different cube sizes and different selection radius to create cognitive conflict.

### Hand movement velocity and its effect on PEN and Pe

The experiment successfully generated two distinct hand movement velocity profiles for the big and small cubes as expected. It was also found that the hand-movement trajectory followed a specific pattern while touching the cube. The participants tended to initially accelerate their hand until they reached the peak velocity, followed by a deceleration before touching the cube. This specific pattern concurs with Meyer et al. (1988) OIP model. The OIP model suggests that when interacting with a 3D object, the hand movement trajectory consists of a ballistic phase followed by a corrective phase divided by the peak velocity.

Interestingly, the conflict trials for the big cubes only seemed to have a ballistic phase without any corrective phase after the peak of the movement velocity. The absence of a corrective phase can be explained by the selection radius of the big cubes that generated premature touch feedback, creating cognitive conflict. An already big cube with an additional large selection radius does not leave enough space for extended hand movements. As such, the feedback was given already during or at the end of the ballistic phase of the movement, and no corrective phase of hand movements was initiated. That could also be the explanation of why there was higher theta power for trials with big cubes compared to small cubes in conflict trials. According to Kalfaoğlu, Stafford, and Milne (2018), uncorrected errors result in stronger modulations of theta power compared to corrected errors. Therefore, our findings of higher theta power coupled with the absence of a corrective movement phase, i.e., uncorrected errors, support their argument.

The results from the conflict trials with big cubes showed a significant difference in higher theta and alpha powers just after the color change feedback, unlike the small cubes. The difference appeared within 50ms after the onset of feedback, which meant the participants did not have a chance to correct their movements. However, the participants were more careful with the small cubes from the outset, requiring fewer adjustments in the corrective stage. This again serves as proof that the OIP model holds in the 3D object selection tasks (Ferran Argelaguet & Andujar, 2013; Meyer et al., 1988).

The hand movement trajectories profiles also affected the PEN and Pe amplitudes. The PEN amplitude in the big cubes was significantly higher than that of the small cubes in conflict condition. We believe it was because, in the big cube for conflict conditions, PENs were evoked during the ballistic phase. The OIP model suggests that the participant would expect less correction and adjustment for an easy target, i.e., big cube, during the ballistic phase. When the cognitive conflict arises, i.e., the big cube changed its color prematurely during the ballistic phase, the participants would need to allocate more cognitive resources than expected to respond to the prediction error and thus larger corresponding PENs. For small cubes in conflict conditions, PENs were evoked during the corrective phase. In the corrective phase, the participants were already engaged in the process of correction and adjustment. The prediction error would require less additional cognitive resources and, thus, a smaller PEN amplitude. In contrast, there were no significant differences in Pe between big and small cubes for conflict conditions. This may be because the Pe amplitude was modulated by the visual stimuli, i.e., change of cube color, which was identical in both conditions (Polich, 2007).

### Origin of cognitive conflict, event-related spectral perturbation, and its correlation with the hand’s velocity

Several past studies have demonstrated that the cognitive conflict originates in ACC and is also a contributor to the family of event-related errors that includes PEN (Carter et al., 1998; Devinsky, Morrell, & Vogt, 1995; van Veen, Cohen, Botvinick, Stenger, & Carter, 2001). Our tests that localize EEG data with ICA (Makeig, Bell, Jung, & Sejnowski, 1996) and dipole fit (Scherg, 1990) concur with those findings. ACC was found to be activated in most participants as they performed the 3D object selection task (see Figure 10). This strengthens proof that ACC is indeed involved in the cognitive process. ACC is the vital hub behind our ability to handle situations of cognitive conflict (Carter et al., 1998; Umemoto, Inzlicht, & Holroyd, 2018). ACC is also known to interact with motor controls in a bottom-up fashion (Rauss & Pourtois, 2013). That cognitive conflict originates in ACC is also aligned with other related tasks, such as action monitoring (Botvinick, Braver, Barch, Carter, & Cohen, 2001), observational errors (van Schie et al., 2004), and prediction errors (Gehrke et al., 2019; Ozkan & Pezzetta, 2018; A. K. Singh et al., 2020; Singh et al., 2018).

To further verify the origin of cognitive conflict, we looked at ERSP in ACC. The results show that frontal theta and alpha power are modulated by cognitive conflict responses in the participants. Modulations in theta power accord with existing theories of frontal theta power variations during tasks that involve cognitive conflict (Arrighi et al., 2016; Zhang, Watrous, Patel, & Jacobs, 2018). However, we founded one difference. Our results showed that theta power decreases in situations of cognitive conflict while other results show an increase – assumed to be the result of phase resetting over a sudden change in behavior, like cognitive conflict, which generates an error-related negativity (Luu, Tucker, & Makeig, 2004; Sauseng, Griesmayr, Freunberger, & Klimesch, 2010). We find evidence to dispute this claim in that phase resetting does not always generate ERN (Yeung, Bogacz, Holroyd, Nieuwenhuis, & Cohen, 2007). Hence, our result supports the Yeung et al. (2007) theory. It seems that the PEN evoked in our experiments was not the result of phase resetting in theta given conflict conditions. Markedly, such arguments further raise a question about why theta power phase resetting did not occur. This requires further experimentation and investigation.

In addition to the theta power modulations discussed in the previous section, modulations in alpha power are also known to be related to errors (van Driel, Ridderinkhof, & Cohen, 2012). Several previous works suggest that the alpha power modulation could be the results of attention and perception (Den Ouden, Kok, & De Lange, 2012), self-awareness (Devinsky et al., 1995), and the observer’s relationship with the person performing the task (Kang, Hirsh, & Chasteen, 2010). We found significantly more alpha power modulation in the conflict trials with the small cubes than the big cubes. This is potentially due to the higher attention requirements from the very beginning of hand movement, which is also in line with attentional process theory Pfurtscheller (2003). Interestingly, the small cubes in the no-conflict trials also showed a significant correlation between the ballistic phase and alpha and beta power. Alpha and beta power are often associated with focused attention and motor inhibition (Foxe & Snyder, 2011; Horimoto, Inagaki, Yano, Sata, & Kaga, 2002; Neuper & Pfurtscheller, 2001), which were both required for our 3D object selection task. Nevertheless, there is still the question of why the same is not true for the conflict trials. The reason could be the dominance of theta power where cognitive conflict exists, which might dissipate the effects of other power bands.

We also found a correlation between delta power and the ballistic phase in the conflict trials with big cubes. The delta power band plays an essential role in the inhibition process, such as cognitive conflict (Harmony, 2013). Due to the lack of a corrective phase, the participants may have needed to inhibit their actions over quite a short period, which would explain such a correlation. In addition to the above, delta band modulation also found in other reaching out and imagined hand-movement tasks (Korik, Sosnik, Siddique, & Coyle, 2018; Zeng et al., 2019).

### Modeling the relationship between PEN, Pe and hand movement velocity, task completion time, and IPQ scores

It is evident so far that the ballistics and corrective phase of hand movements have an impact on PEN and Pe amplitude in conflict trials. It was found that the ballistic phase for small cube modulated the PEN but slightly the Pe components of the ERP. Further, the ballistic of hand movement velocity with the behavior information from the IPQ (QVR and SEVR) was able to account for more than 48% of the variability in PEN for both small and big cubes. However, Pe is only a predictor with big cubes and even then, only accounts for 28% of the variability. We attribute this weak correlation to the hand movement trajectory. As mentioned before, the hand movements with big cubes had a ballistic phase of hand movement only, which was predominantly guided by combined proprioceptive and visual sensory information as feedback. Therefore, ballistic of hand movement velocity alone was able to predict the PEN’s amplitude. Although, including the realism and PAVR IPQ scores, along with the task completion time, increased the ability to predict PEN by more than 14%.

Interestingly, the IPQ score seems to play an essential role in modeling PEN and Pe. In past studies, the participants’ experience was found to be highly related to their interactions with the environment and how the environment affects behavior (Balconi & Crivelli, 2010; Devinsky et al., 1995). A previous experiment by Singh et al. (2018) also shows that visual appearance affects cognitive conflict and is related to both the level of realism (F. Argelaguet, Hoyet, Trico, & Lecuyer, 2016) and the behavior inhibition score (Carver & White, 1994). Our findings are in-line with these studies and explain why the IPQ scores from the participants played such an influential role in predicting PEN and Pe with the ballistic phase and task completion time. The participant’s interactive experience with VR, such as their control over the scene, its realism, etc. made it easier to translate their feelings toward the cognitive conflict. The participants with higher experience with VR also demonstrated higher PEN amplitudes in the cognitive conflict conditions.

Overall, the results indicate that hand movement velocity in an object selection task plays an essential role in handling cognitive conflicts, most likely because movement-related proprioception is critical for corrective hand movements (Bagesteiro, Sarlegna, & Sainburg, 2005). The results from behavior, EEG, regression, and correlation support the conclusion that the velocity of the hand’s movement impacts cognitive conflict processing in 3D object selection tasks. Such a finding is only possible due to the nature of the task. This task is one of the first of its kind to involve active motor control in the field of neuroscience, falling into the category of mobile-brain/body imaging (MoBI) (Gramann et al., 2011; Gramann, Jung, Ferris, Lin, & Makeig, 2014; Makeig, Gramann, Jung, Sejnowski, & Poizner, 2009). This finding has implications for our understanding of how proprioceptive and visual sensory information are integrated and together work toward cognitive control. These findings would be beneficial for enhancing user experiences in real and virtual environments with an adaptive system for therapeutic and entertainment purposes.

## Conclusion

In this study, we investigated the impact of hand movement velocity on cognitive conflict processing. We designed an experimental scenario to invoke different velocity profiles during 3D object selection. The participants were asked to grasp virtual cubes in a series of 2×2 factor trials: the first condition being the size of the cube – big or small; the second being the selection radius of the cube to induce a conflict/non-conflict situation. The results of regression analysis with PEN, Pe, and the participants’ IPQ scores show that PEN is modulated in the ballistic phase and highly related to proprioceptive information. Additionally, previous experience with VR technology, as self-reported in the IPQ, also significantly impacts cognitive processing.

## Footnotes

2 https://www.unity.com/

3 http://www.talairach.org/

## Notes

### Competing Interest Statement

The authors have declared no competing interest.

## References

Argelaguet, F., & Andujar, C. (2013). A survey of 3D object selection techniques for virtual environments. Computers & Graphics, 37(3), 121–136. doi:https://doi.org/10.1016/j.cag.2012.12.003

Argelaguet, F., Hoyet, L., Trico, M., & Lecuyer, A. (2016, 19–23 March 2016). The role of interaction in virtual embodiment: Effects of the virtual hand representation. Paper presented at the 2016 IEEE Virtual Reality (VR).

Arrighi, P., Bonfiglio, L., Minichilli, F., Cantore, N., Carboncini, M. C., Piccotti, E.,… Andre, P. (2016). EEG Theta Dynamics within Frontal and Parietal Cortices for Error Processing during Reaching Movements in a Prism Adaptation Study Altering Visuo-Motor Predictive Planning. PLoS One, 11(3), e0150265. doi:10.1371/journal.pone.0150265

Bagesteiro, L. B., Sarlegna, F. R., & Sainburg, R. L. (2005). Differential influence of vision and proprioception on control of movement distance. Experimental Brain Research, 171(3), 358. doi:10.1007/s00221-005-0272-y

Balconi, M., & Crivelli, D. (2010). FRN and P300 ERP effect modulation in response to feedback sensitivity: The contribution of punishment-reward system (BIS/BAS) and Behaviour Identification of action. Neuroscience Research, 66(2), 162–172. doi:https://doi.org/10.1016/j.neures.2009.10.011

Bedikian, R. (2013). Understanding Latency: Part 1. Online Retrieved from http://blog.leapmotion.com/understanding-latency-part-1/

Botvinick, M. M., Braver, T. S., Barch, D. M., Carter, C. S., & Cohen, J. D. (2001). Conflict monitoring and cognitive control. Psychol Rev, 108(3), 624–652.

Carter, C. S., Braver, T. S., Barch, D. M., Botvinick, M. M., Noll, D., & Cohen, J. D. (1998). Anterior cingulate cortex, error detection, and the online monitoring of performance. Science, 280(5364), 747–749.

Carver, C. S., & White, T. L. (1994). Behavioral inhibition, behavioral activation, and affective responses to impending reward and punishment: The BIS/BAS Scales. Journal of personality and social psychology, 67(2), 319–333. doi:10.1037/0022-3514.67.2.319

Chatrian, G. E., Lettich, E., & Nelson, P. L. (1985). Ten Percent Electrode System for Topographic Studies of Spontaneous and Evoked EEG Activities. American Journal of EEG Technology, 25(2), 83–92. doi:10.1080/00029238.1985.11080163

Chaumon, M., Bishop, D. V. M., & Busch, N. A. (2015). A practical guide to the selection of independent components of the electroencephalogram for artifact correction. Journal of Neuroscience Methods, 250(Supplement C), 47–63. doi:https://doi.org/10.1016/j.jneumeth.2015.02.025

Coull, J. T., & Nobre, A. C. (1998). Where and when to pay attention: the neural systems for directing attention to spatial locations and to time intervals as revealed by both PET and fMRI. J Neurosci, 18(18), 7426–7435.

Delorme, A., & Makeig, S. (2004). EEGLAB: an open source toolbox for analysis of single-trial EEG dynamics including independent component analysis. Journal of Neuroscience Methods, 134(1), 9–21. doi:http://dx.doi.org/10.1016/j.jneumeth.2003.10.009

Den Ouden, H., Kok, P., & De Lange, F. (2012). How Prediction Errors Shape Perception, Attention, and Motivation. Frontiers in Psychology, 3(548). doi:10.3389/fpsyg.2012.00548

Desmurget, M., & Grafton, S. (2000). Forward modeling allows feedback control for fast reaching movements. Trends Cogn Sci, 4(11), 423–431. doi:10.1016/s1364-6613(00)01537-0

Devinsky, O., Morrell, M. J., & Vogt, B. A. (1995). Contributions of anterior cingulate cortex to behaviour. Brain, 118(1), 279–306. doi:10.1093/brain/118.1.279

Donchin, E., & Coles, M. G. H. (2010). Is the P300 component a manifestation of context updating? Behavioral and Brain Sciences, 11(3), 357–374. doi:10.1017/S0140525X00058027

Eriksen, B. A., & Eriksen, C. W. (1974). Effects of noise letters upon the identification of a target letter in a nonsearch task. Perception & Psychophysics, 16(1), 143–149. doi:10.3758/bf03203267

Falkenstein, M., Hohnsbein, J., Hoormann, J., & Blanke, L. (1991). Effects of crossmodal divided attention on late ERP components. II. Error processing in choice reaction tasks. Electroencephalogr Clin Neurophysiol, 78(6), 447–455.

Fan, J., Flombaum, J. I., McCandliss, B. D., Thomas, K. M., & Posner, M. I. (2003). Cognitive and brain consequences of conflict. Neuroimage, 18(1), 42–57.

Foxe, J. J., & Snyder, A. C. (2011). The Role of Alpha-Band Brain Oscillations as a Sensory Suppression Mechanism during Selective Attention. Frontiers in Psychology, 2, 154. doi:10.3389/fpsyg.2011.00154

Gehring, W. J., Goss, B., Coles, M. G. H., Meyer, D. E., & Donchin, E. (1993). A Neural System for Error Detection and Compensation. Psychological Science, 4(6), 385–390. Retrieved from http://www.jstor.org/stable/40062567

Gehrke, L., Akman, S., Lopes, P., Chen, A., Singh, A. K., Chen, H.-T., Gramann, K. (2019). Detecting Visuo-Haptic Mismatches in Virtual Reality using the Prediction Error Negativity of Event-Related Brain Potentials. Paper presented at the Proceedings of the 2019 CHI Conference on Human Factors in Computing Systems, Glasgow, Scotland Uk.

Gramann, K., Gwin, J. T., Ferris, D. P., Oie, K., Jung, T.-P., Lin, C.-T., Makeig, S. (2011). Cognition in action: imaging brain/body dynamics in mobile humans. Rev Neurosci, 22(6), 593–608.

Gramann, K., Jung, T.-P., Ferris, D. P., Lin, C.-T., & Makeig, S. (2014). Toward a new cognitive neuroscience: modeling natural brain dynamics. Frontiers in Human Neuroscience, 8, 444.

Green, S. B., Salkind, N. J., & Akey, T. M. (1997). Using SPSS for Windows; analyzing and understanding data: Prentice Hall PTR.

Halgren, E., Marinkovic, K., & Chauvel, P. (1998). Generators of the late cognitive potentials in auditory and visual oddball tasks. Electroencephalogr Clin Neurophysiol, 106(2), 156–164. doi:https://doi.org/10.1016/S0013-4694(97)00119-3

Harmony, T. (2013). The functional significance of delta oscillations in cognitive processing. Frontiers in Integrative Neuroscience, 7, 83–83. doi:10.3389/fnint.2013.00083

Holroyd, C. B., & Coles, M. G. H. (2002). The neural basis of human error processing: reinforcement learning, dopamine, and the error-related negativity. Psychol Rev, 109(4), 679–709. doi:10.1037/0033-295x.109.4.679

Horimoto, R., Inagaki, M., Yano, T., Sata, Y., & Kaga, M. (2002). Mismatch negativity of the color modality during a selective attention task to auditory stimuli in children with mental retardation. Brain Dev, 24(7), 703–709.

Jungnickel, E., & Gramann, K. (2016). Mobile Brain/Body Imaging (MoBI) of Physical Interaction with Dynamically Moving Objects. Frontiers in Human Neuroscience, 10, 306. doi:10.3389/fnhum.2016.00306

Kalfaoğlu, Ç., Stafford, T., & Milne, E. (2018). Frontal theta band oscillations predict error correction and posterror slowing in typing. Journal of Experimental Psychology: Human Perception and Performance, 44(1), 69–88. doi:10.1037/xhp0000417

Kang, S. K., Hirsh, J. B., & Chasteen, A. L. (2010). Your mistakes are mine: Self-other overlap predicts neural response to observed errors. Journal of Experimental Social Psychology, 46(1), 229–232. doi:https://doi.org/10.1016/j.jesp.2009.09.012

Kopp, B., Rist, F., & Mattler, U. W. E. (1996). N200 in the flanker task as a neurobehavioral tool for investigating executive control. Psychophysiology, 33(3), 282–294. doi:10.1111/j.1469-8986.1996.tb00425.x

Korik, A., Sosnik, R., Siddique, N., & Coyle, D. (2018). Decoding Imagined 3D Hand Movement Trajectories From EEG: Evidence to Support the Use of Mu, Beta, and Low Gamma Oscillations. Front Neurosci, 12(130). doi:10.3389/fnins.2018.00130

Luu, P., Tucker, D. M., & Makeig, S. (2004). Frontal midline theta and the error-related negativity: neurophysiological mechanisms of action regulation. Clinical Neurophysiology, 115(8), 1821–1835. doi:https://doi.org/10.1016/j.clinph.2004.03.031

Makeig, S., Bell, A. J., Jung, T.-P., & Sejnowski, T. J. (1996). Independent component analysis of electroencephalographic data. Paper presented at the Advances in neural information processing systems.

Makeig, S., Gramann, K., Jung, T.-P., Sejnowski, T. J., & Poizner, H. (2009). Linking brain, mind and behavior. International Journal of Psychophysiology, 73(2), 95–100.

Meyer, D. E., Abrams, R. A., Kornblum, S., Wright, C. E., & Smith, J. E. (1988). Optimality in human motor performance: ideal control of rapid aimed movements. Psychol Rev, 95(3), 340–370.

Mognon, A., Jovicich, J., Bruzzone, L., & Buiatti, M. (2011). ADJUST: An automatic EEG artifact detector based on the joint use of spatial and temporal features. Psychophysiology, 48(2), 229–240. doi:doi:10.1111/j.1469-8986.2010.01061.x

Montgomery, E. B., Huang, H., & Assadi, A. (2005). Unsupervised clustering algorithm for N-dimensional data. Journal of Neuroscience Methods, 144(1), 19–24. doi:https://doi.org/10.1016/j.jneumeth.2004.10.015

Neuper, C., & Pfurtscheller, G. (2001). Event-related dynamics of cortical rhythms: frequency-specific features and functional correlates. Int J Psychophysiol, 43(1), 41–58.

Ozkan, D. G., & Pezzetta, R. (2018). Predictive monitoring of actions, EEG recordings in virtual reality. J Neurophysiol, 119(4), 1254–1256. doi:10.1152/jn.00825.2017

Padrao, G., Gonzalez-Franco, M., Sanchez-Vives, M. V., Slater, M., & Rodriguez-Fornells, A. (2016). Violating body movement semantics: Neural signatures of self-generated and external-generated errors. Neuroimage, 124(Pt A), 147–156. doi:10.1016/j.neuroimage.2015.08.022

Padrao, G., Rodriguez-Herreros, B., Perez Zapata, L., & Rodriguez-Fornells, A. (2015). Exogenous capture of medial-frontal oscillatory mechanisms by unattended conflicting information. Neuropsychologia, 75, 458–468. doi:10.1016/j.neuropsychologia.2015.07.004

Pfurtscheller, G. (2003). Induced Oscillations in the Alpha Band: Functional Meaning. Epilepsia, 44, 2–8. doi:10.1111/j.0013-9580.2003.12001.x

Polich, J. (2007). Updating P300: An integrative theory of P3a and P3b. Clinical Neurophysiology, 118(10), 2128–2148. doi:https://doi.org/10.1016/j.clinph.2007.04.019

Rauss, K., & Pourtois, G. (2013). What is Bottom-Up and What is Top-Down in Predictive Coding? Frontiers in Psychology, 4(276). doi:10.3389/fpsyg.2013.00276

Sauseng, P., Griesmayr, B., Freunberger, R., & Klimesch, W. (2010). Control mechanisms in working memory: A possible function of EEG theta oscillations. Neuroscience & Biobehavioral Reviews, 34(7), 1015–1022. doi:http://dx.doi.org/10.1016/j.neubiorev.2009.12.006

Scheidt, R. A., Conditt, M. A., Secco, E. L., & Mussa-Ivaldi, F. A. (2005). Interaction of Visual and Proprioceptive Feedback During Adaptation of Human Reaching Movements. J Neurophysiol, 93(6), 3200–3213. doi:10.1152/jn.00947.2004

Scherg, M. (1990). Fundamentals of dipole source potential analysis. Auditory evoked magnetic fields and electric potentials. Advances in audiology, 6, 40–69.

Schlüter, C., Fraenz, C., Pinnow, M., Friedrich, P., Güntürkün, O., & Genç, E. (2018). The Structural and Functional Signature of Action Control. Psychological Science, 29(10), 1620–1630. doi:10.1177/0956797618779380

Schubert, T. W. (2003). The sense of presence in virtual environments: A three-component scale measuring spatial presence, involvement, and realness. Zeitschrift für Medienpsychologie, 15(2), 69–71.

Singh, A. K., Chen, H.-T., Gramann, K., & Lin, C.-T. (2020). Intra-individual Completion Time Modulates the Prediction Error Negativity in a Virtual 3D Object Selection Task. IEEE Transactions on Cognitive and Developmental Systems.

Singh, A. K., Chen, H., Gramann, K., & Lin, C. (2020). Intraindividual Completion Time Modulates the Prediction Error Negativity in a Virtual 3-D Object Selection Task. IEEE Transactions on Cognitive and Developmental Systems, 12(2), 354–360. doi:10.1109/TCDS.2020.2991301

Singh, A. K., Chen, H. T., Cheng, Y. F., King, J. T., Ko, L. W., Gramann, K., & Lin, C. T. (2018). Visual Appearance Modulates Prediction Error in Virtual Reality. IEEE Access, 6, 24617–24624. doi:10.1109/ACCESS.2018.2832089

Soukoreff, R. W., & MacKenzie, I. S. (2004). Towards a standard for pointing device evaluation, perspectives on 27 years of Fitts’ law research in HCI. International Journal of Human-Computer Studies, 61(6), 751–789.

Squires, N. K., Squires, K. C., & Hillyard, S. A. (1975). Two varieties of long-latency positive waves evoked by unpredictable auditory stimuli in man. Electroencephalogr Clin Neurophysiol, 38(4), 387–401.

Stroop, J. R. (1935). Studies of interference in serial verbal reaction. J Exp Psychol, 18. doi:10.1037/h0054651

Umemoto, A., Inzlicht, M., & Holroyd, C. B. (2018). Electrophysiological indices of anterior cingulate cortex function reveal changing levels of cognitive effort and reward valuation that sustain task performance. Neuropsychologia. doi:https://doi.org/10.1016/j.neuropsychologia.2018.06.010

van Driel, J., Ridderinkhof, K. R., & Cohen, M. X. (2012). Not All Errors Are Alike: Theta and Alpha EEG Dynamics Relate to Differences in Error-Processing Dynamics. The Journal of Neuroscience, 32(47), 16795. Retrieved from http://www.jneurosci.org/content/32/47/16795.abstract

van Schie, H. T., Mars, R. B., Coles, M. G., & Bekkering, H. (2004). Modulation of activity in medial frontal and motor cortices during error observation. Nat Neurosci, 7(5), 549–554. doi:10.1038/nn1239

van Veen, V., Cohen, J. D., Botvinick, M. M., Stenger, V. A., & Carter, C. S. (2001). Anterior cingulate cortex, conflict monitoring, and levels of processing. Neuroimage, 14(6), 1302–1308. doi:10.1006/nimg.2001.0923

Weichert, F., Bachmann, D., Rudak, B., & Fisseler, D. (2013). Analysis of the Accuracy and Robustness of the Leap Motion Controller. Sensors (Basel, Switzerland), 13(5), 6380–6393. doi:10.3390/s130506380

West, R., & Alain, C. (1999). Event-related neural activity associated with the Stroop task. Cognitive Brain Research, 8(2), 157–164. doi:https://doi.org/10.1016/S0926-6410(99)00017-8

Yeung, N., Bogacz, R., Holroyd, C. B., Nieuwenhuis, S., & Cohen, J. D. (2007). Theta phase resetting and the error-related negativity. Psychophysiology, 44(1), 39–49. doi:doi:10.1111/j.1469-8986.2006.00482.x

Zeng, H., Sun, Y., Xu, G., Wu, C., Song, A., Xu, B.,… Hu, C. (2019). The Advantage of Low-Delta Electroencephalogram Phase Feature for Reconstructing the Center-Out Reaching Hand Movements. Front Neurosci, 13(480). doi:10.3389/fnins.2019.00480

Zhang, H., Watrous, A. J., Patel, A., & Jacobs, J. (2018). Theta and Alpha Oscillations Are Traveling Waves in the Human Neocortex. Neuron, 98(6), 1269–1281.e1264. doi:10.1016/j.neuron.2018.05.019

